# Differential excretory/secretory proteome of the adult female and male stages of the human blood fluke, *Schistosoma mansoni*

**DOI:** 10.1101/2022.05.22.492965

**Authors:** Eric T. Kenney, Victoria H. Mann, Wannaporn Ittiprasert, Bruce A. Rosa, Makedonka Mitreva, Bethany K. Bracken, Alex Loukas, Paul J. Brindley, Javier Sotillo

## Abstract

Intricate molecular communication between the schistosome (flatworms) and its mammalian host, as well as between paired male and female schistosomes has shaped the secreted proteome of these flatworms. Whereas the schistosome egg is responsible for the disease manifestations of chronic schistosomiasis, the long lived, adult female and male stages also release mediators that facilitate their long-lived intra-vascular existence in a hostile niche where they are bathed in immune cells and effector molecules. However, despite their importance, no studies have focused on analysing the excretory/secretory products (ESPs) from adult schistosomes.

Herein, ESPs from cultured *Schistosoma mansoni* male or female adult worms were identified, quantified, compared and contrasted using a label-free proteomic approach. Approximately 1,000 proteins were identified, from which almost 800 could be finally quantified. Considering the proteins uniquely identified and proteins with a significantly regulated expression pattern in male or female flukes, a total of 370 and 140 proteins were more abundantly secreted by males and females, respectively. Using functional analysis networks showing the gene ontology terms and KEGG pathways with the highest significance, we observed that male schistosomes secrete proteins related to carbohydrate metabolism, cytoskeletal organisation more abundantly than females, while female worms secreted more hydrolases and proteins involved in cellular homeostasis than males.

This analysis doubles the number of previously reported ESPs from *S. mansoni*, contributing to a better understanding of the host-parasite dynamic interactions. Furthermore, these findings expand potential vaccine and diagnostic candidates for this neglected tropical disease pathogen, which will enable deeper understanding of the molecular communication critical to parasitism.

## 1 Introduction

Schistosomiasis is a major neglected tropical disease and is considered the most important helminthiasis in terms of morbidity and mortality. More than 200 million people are infected worldwide with 700 million at risk of infection. This remains a major public health problem, particularly in sub-Saharan Africa (Colley et al., 2014;McManus et al., 2018). Human schistosomiasis is caused by six species of blood flukes: *Schistosoma guineensis, S. haematobium, S. intercalatum, S. japonicum, S. mansoni*, and *S. mekongi*. Nowadays, the predominant human species are *S. haematobium* and *S. mansoni*, given the reduction of infection in recent decades caused by *S. japonicum* in the Yangtze River basin provinces of China (Wang et al., 2021). Urogenital schistosomiasis caused by hybrids of *S. haematobium* and *S. bovis* and relatives is spreading in West Africa (Webster et al., 2006;Huyse et al., 2009) and in Corsica (Boissier et al., 2016;Rothe et al., 2021).

Male and female schistosomes dwell in copula within the mesenteric veins (*S. mansoni, S. japonicum*) or the vesical venous plexus (*S. haematobium*) of the human, laying hundreds to thousands (depending on the species) of fertilized eggs each day. The eggs traverse the intestinal wall (e.g., *S. mansoni*) or the bladder wall (*S. haematobium*) and exit the host to the external environment in feces or urine, respectively. However, many eggs fail to exit the infected person and are retained in host tissues where they induce inflammation, granuloma, and fibrosis (McManus et al., 2018). In the external environment, the eggs hatch when they reach freshwater, each releasing a free-living larva, the miracidium, which is ciliated and seeks to infect the obligate intermediate host, a snail to continue the transmission of the disease. Within the snail, the schistosome undergoes cycles of asexual reproduction through mother and daughter sporocyst stages, eventually shedding thousands of cercariae into the water. The cycles of asexual reproduction within the snail require several weeks before cercariae are released. The cercaria is the infectious developmental stage for humans and other permissive mammals. After penetrating the skin, the cercaria sheds its tail and the juvenile larva, the schistosomulum, migrates within the circulatory system, reaching the lungs, the liver, and eventually the portal venous system or the venous system that drains the pelvic organs where the fully mature flukes pair and the female produces eggs, completing the developmental cycle.

Whereas the schistosome egg is responsible for the disease manifestations of chronic schistosomiasis, as well as orchestrating the hallmark immunological transition from a Th1 to Th2 response (Pearce and MacDonald, 2002;Schwartz and Fallon, 2018;Acharya et al., 2021), the long lived, adult female and male stages also release mediators that facilitate their long-lived intra-vascular existence in a hostile niche where they are bathed in immune cells and effector molecules. These mediators, also known as excretory/secretory products (ESPs), are secreted (or released) from the esophageal gland, the gut epithelium and from the tegument of schistosomes, making it a highly diverse mixture of molecules. ESPs from several developmental stages and species of schistosomes have been described in depth, e.g., (Liu et al., 2009;Mathieson and Wilson, 2010;Hall et al., 2011;Dvořák et al., 2016;Sotillo et al., 2016;Floudas et al., 2017;De Marco Verissimo et al., 2019;Sotillo et al., 2019;Kifle et al., 2020;Neves et al., 2020;Chen et al., 2022), although the diversity, role, and packaging of these secreted and excreted antigens, including as cargo within extracellular vesicles, may not yet be fully characterized or deciphered, e.g., (Acharya et al., 2021). An early study of adults stage *S. japonicum* ESPs showed the presence of canonical proteins such as metabolic enzymes, heat shock proteins (HSPs), detoxification proteins, and peptidases (Liu et al., 2009). Other studies focusing strictly on the schistosome “vomitus” (proteins secreted only by the gut epithelium) highlighted the presence of different saposins, ferritins, and cathepsins among other molecules (Hall et al., 2011). Furthermore, tetraspanins, annexins, calpain, and several transporters and cytoskeletal proteins have been identified on the tegument of *S. mansoni* adult worms by different techniques (Braschi et al., 2006a;Braschi et al., 2006b;Braschi and Wilson, 2006).

Here, the ESPs in cultured supernatants from adult male and adult female *S. mansoni* were isolated and compared using a label-free proteomic approach for the first time, increasing the coverage of the published secretome. About 1,000 proteins were identified, from which ∼ 800 could be finally quantified. In sum, this new analysis at least doubles the number of proteins known in these extracts (Hall et al., 2011;Wilson, 2012), substantially expanding the catalogue of ESPs from *S. mansoni*, which provides new insights of the host-parasite interplay. In turn, we augment the number of potential vaccine and diagnostic candidates listed previously for this neglected tropical disease agent.

## 2 Materials and methods

### 2.1 Ethics

Mice experimentally infected with *S. mansoni*, obtained from the Schistosomiasis Resource Center (SRC) at the Biomedical Research Institute (BRI), MD were housed at the Animal Research Facility of the George Washington University (GWU), which is accredited by the American Association for Accreditation of Laboratory Animal Care (AAALAC no. 000347) and has an Animal Welfare Assurance on file with the National Institutes of Health, Office of Laboratory Animal Welfare, OLAW assurance number A3205-01. All procedures employed were consistent with the Guide for the Care and Use of Laboratory Animals. The Institutional Animal Care and Use Committee (IACUC) at GWU approved the protocol used for maintenance of mice and recovery of schistosomes.

### 2.2 Schistosomes

Swiss-Webster albino mice were euthanized seven weeks after infection with *S. mansoni*, livers were removed at necropsy, schistosome eggs recovered from the livers, and adult worms from the portal circulation as described (Dalton et al., 1997).

### 2.3 Isolation of adult excretory/secretory products

For the collection of excretory-secretory products (ESPs), adult *S. mansoni* were provided by Schistosomiasis Resource Center (SRC) of the Biomedical Research Institute (BRI), Rockville, MD. The worms were sorted with forceps to separate males and females, rinsed briefly in 1x phosphate buffered saline (PBS) (Corning), and subsequently transferred to 100 × 20 mm tissue culture plates (Sarstedt) containing 30 mL of serum-free Dulbecco’s Modification of Eagle’s Medium (DMEM) (Corning) supplemented with 2% Antibiotic-Antimycotic (Gibco). Secretion of ESP into the serum-free medium was facilitated by continuous incubation at 37°C, 5% CO_2_ in air (Neves et al., 2020). At intervals of 24 hours over seven days, 20 mL of culture supernatant was removed with minimal disturbance to the schistosomes and stored at -80°C. The drawn medium was retained for storage and was replaced with fresh medium to the culture at each time point. At the conclusion of the collection period, the ESP-containing media were thawed gradually on wet ice, after which ESP was concentrated using Centricon Plus-70 Centrifugal Filter Units (Millipore) featuring a 3 kDa nominal molecular size limit. Concentration by centrifugation on the 3 kDa membrane was undertaken at 3,220 rpm at 4°C using an Eppendorf 5810R centrifuge fitted with an A-4-62 swinging bucket rotor. Concentrated ESP was resuspended and reconcentrated twice using volumes of chilled PBS equivalent to the starting volume of the sample. Protein concentration was ascertained by the Pierce BCA Protein Assay Kit (Thermo Fisher) method, and concentrated ESP was aliquoted and stored at - 80°C.

### 2.4 Mass spectrometry analysis

Three biological replicates of ES from males, females and mixed samples were individually processed as follows. Samples were freeze-dried and dissolved with 22 mL of 50 mM ammonium bicarbonate. Two (2) mL was used to quantify the protein concentration with Qubit (Invitrogen) reagent according to the manufacturer’s instructions. Ten (10) mg of protein was taken and volumes set to 22.5 mL of 50 mM ABC. Reduction and alkylation were performed by incubating samples at 60 °C for 20 min with 2 mM dithiothreitol followed by a 30 min incubation at RT in the dark with 5.5 mM 2-iodoacetamide. Samples were then in-solution digested with 400 ng trypsin overnight at 37 ºC and acidified with 10% TFA to a final concentration of 1%. Digested peptides were finally concentrated by speed vacuum to 15 μL.

Five (5) μl of peptide mixtures were loaded onto a trap column (3μ C18-CL, 350 μm x 0.5mm; Eksigent Technologies, Redwood City, CA) and desalted with 0.1% TFA at 5 μl/min during 5 min. The peptides were then loaded onto an analytical column (3μ C18-CL 120 □, 0.075 × 150 mm; Eksigent) equilibrated in 5% acetonitrile 0.1% FA. Elution was carried out with a linear gradient of 15-40 % B in A for 60 min (A: 0.1% FA; B: ACN, 0.1% FA) at a flow rate of 300 nL/min. Peptides were analysed in a mass spectrometer nanoESI qQTOF (6600+ TripleTOF, ABSCIEX). Sample was ionized in a Source Type: Optiflow < 1 μL Nano applying 3.0 kV to the spray emitter at 175 °C. Analysis was carried out in a data-dependent mode. Survey MS1 scans were acquired from 350–1400 m/z for 250 ms. The quadrupole resolution was set to ‘LOW’ for MS2 experiments, which were acquired 100–1500 m/z for 25 ms in ‘high sensitivity’ mode. The following switch criteria were used: charge: 2+ to 4+; minimum intensity; 250 counts per second (cps). Up to 100 ions were selected for fragmentation after each survey scan. Dynamic exclusion was set to 15 s.

### 2.5 Database search and protein quantification

Database searches were performed using FragPipe (v16.0) with MSFragger (v3.3) (Kong et al., 2017) and Philosopher (v4.0) (da Veiga Leprevost et al., 2020) against a concatenated target/decoy database consisting of the *S. mansoni* proteome (UP000008854) and common contaminants from Uniprot (downloaded 30 June 2021; 14,615 proteins). For the MSFragger analysis, precursor and fragment mass tolerance were both set to 20 ppm. Mass calibration and parameter optimization were enabled, and isotope error was set to 0/1/2 with two missed trypsin cleavages allowed. The peptide length was set from 7 to 50, and the peptide mass was set to 500 to 5000 Da. Carbamidomethylation of C (+57.021464 Da) was set as fixed modification and Oxidation of M (+15.994915 Da) and acetylation of protein N-term (+42.010565 Da) as variable modifications. Philosopher (da Veiga Leprevost et al., 2020) with PeptideProphet (Keller et al., 2002) and ProteinProphet (Nesvizhskii et al., 2003) was used to estimate the identification FDR. The PSMs were filtered at 1% PSM and 1% protein identification FDR. Quantification and match between runs (MBR) was performed with IonQuant using default values (Yu et al., 2021).

Mass spectrometry data along with the identification results have been deposited in the ProteomeXchange Consortium via the PRIDE partner repository (Vizcaino et al., 2013) with the dataset identifier PXD030699.

### 2.6 Bioinformatic analysis of proteomic sequence data

Label-free quantitative (LFQ) analysis of identified proteins was performed with the MSstats R package (Choi et al., 2014) using default parameters (equalizeMedians as normalization method; log transformation: 2; Tukey’s median polish as the summary method; censored values in intensity column: null and MBimpute: false). Using a power calculation of 0.9 and FDR of 0.05, fold-changes were considered as significant when ≥ 2.450 and adjusted *p*-value□≤□0.05. STRINGDB https://string-db.org was used to perform a PPI analysis based on confidence in the interaction (minimum required interaction score ≥ 0.7) and the network was visualized using Cytoscape. The Cytoscape plugin ClueGO was used to integrate the Kyoto Encyclopedia of Genes and Genomes (KEGG), and Gene Ontology information (including biological processes, immune processes and molecular functions) (Bindea et al., 2009). The enrichment tests for terms and groups were two-sided tests based on the hyper-geometric distribution with a Kappa Score Threshold of 0.4. All GO terms that were significant with *P* < 0.05 (after correcting for multiple testing using the Bonferroni step down false discovery rate correction), ranged between 3-8 tree intervals and contained a minimum of three genes (representing at least 4% from the total number of associated genes) were selected for further analysis. For each group/cluster, only the node with the smallest adjusted *P*-value was annotated.

## 3 Results

### 3.1 *S. mansoni* adult males secrete more proteins in culture than adult female worms

*S. mansoni* adult worms were obtained from experimentally infected mice and separated into three different groups: males (M), females (F) or left coupled as male and female pairs (MF). The worms were cultured for seven days and the secreted proteins analysed by LC-MS/MS. A label-free quantitative (LFQ) analysis was performed to identify the proteins with significantly up- or down-regulated abundance in the secretomes of all three sample groups. An input file containing unique and razor peptides for 934 validated proteins was generated by MSFragger (Supplementary Table S1). Data was normalised using medians of summed intensities (Supplementary Figure 1). MSstats was used to estimate the power of the analysis performed. For our analysis, and to have a power calculation = 0.9 and FDR = 0.05, fold-changes were considered as significant when ≥ 2.450 (Log2 ≥ 1.3) and the adjusted P-value□≤□0.05 (Supplementary Figure 2).

The quantitative proteomic analysis revealed a clear sex-dependent protein profile. For the M vs. F comparison, a total of 793 proteins were quantified, from which 237 and 75 were uniquely detected in males and females, respectively (Supplementary Table S2). Furthermore, the relative abundance of 133 proteins was significantly higher in the secretome of males (Table 1; Figure 1; Supplementary Table S2), while the abundance level of 65 proteins was significantly higher in the secretome of females (Table 2; Figure 1; Supplementary Table S2).

**Table 1.**
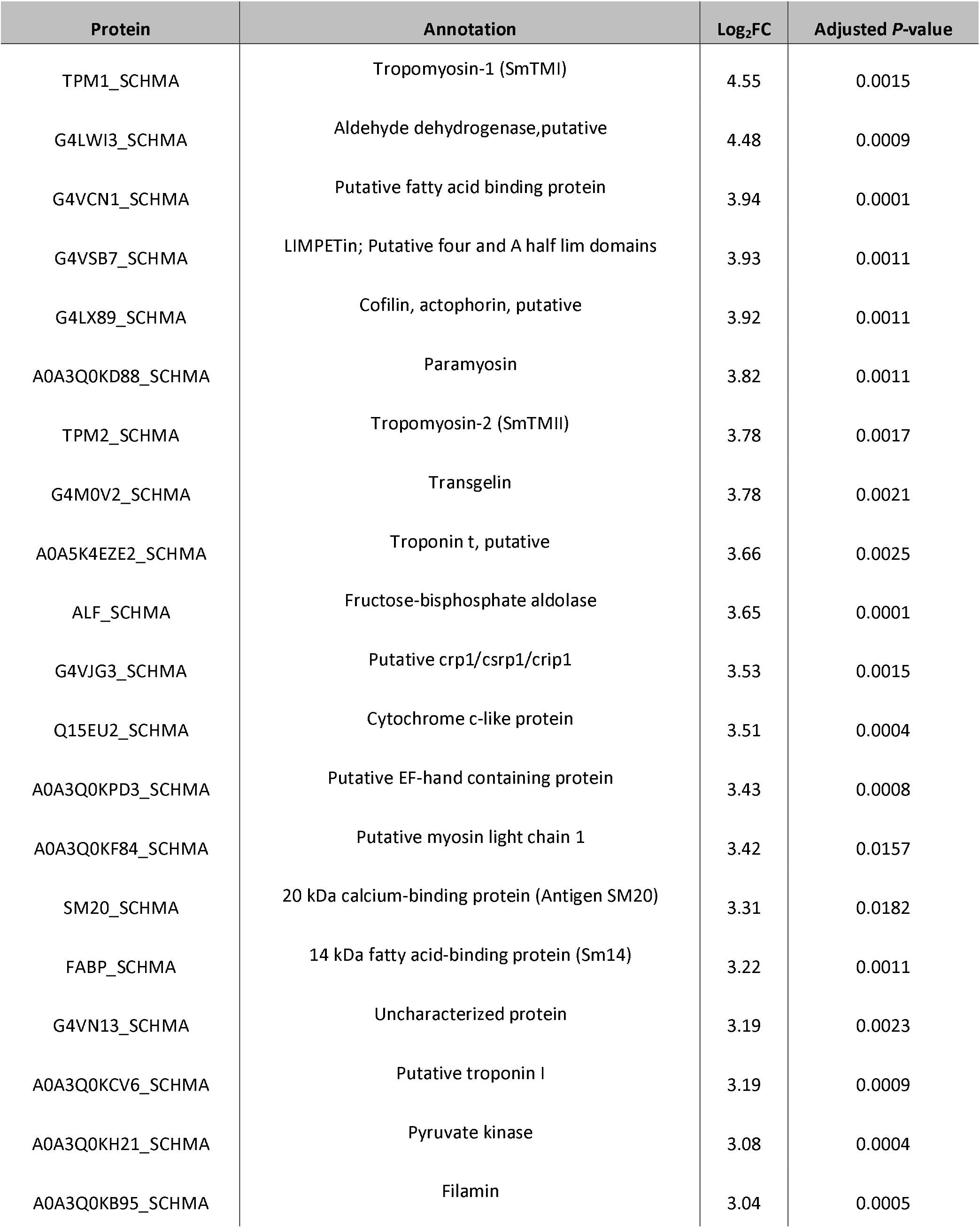
Top 20 most abundant proteins secreted by male *Schistosoma mansoni* adult worms.

**Table 2.**
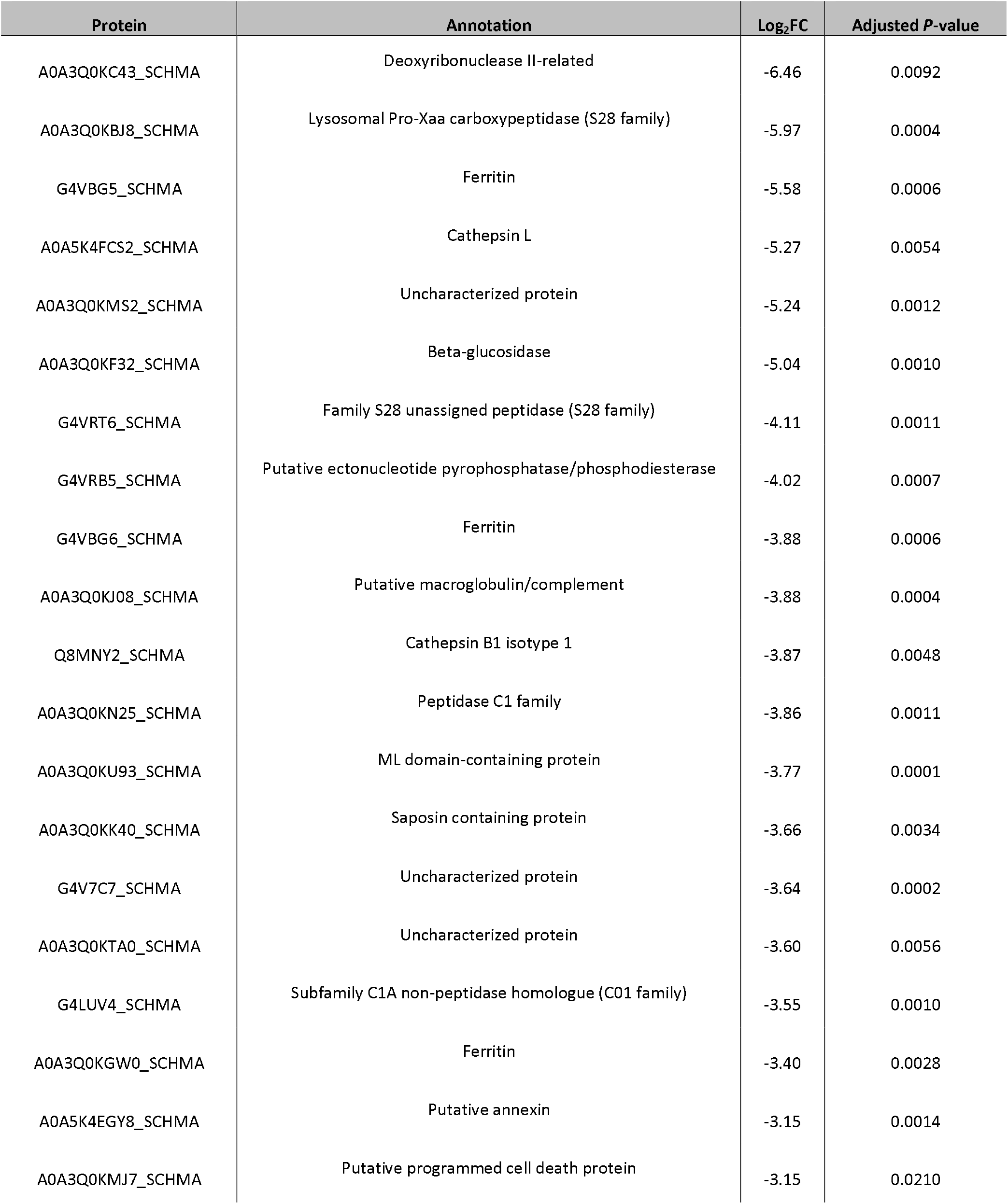
Top 20 most abundant proteins secreted by female *Schistosoma mansoni* adult worms.

**Figure 1.**
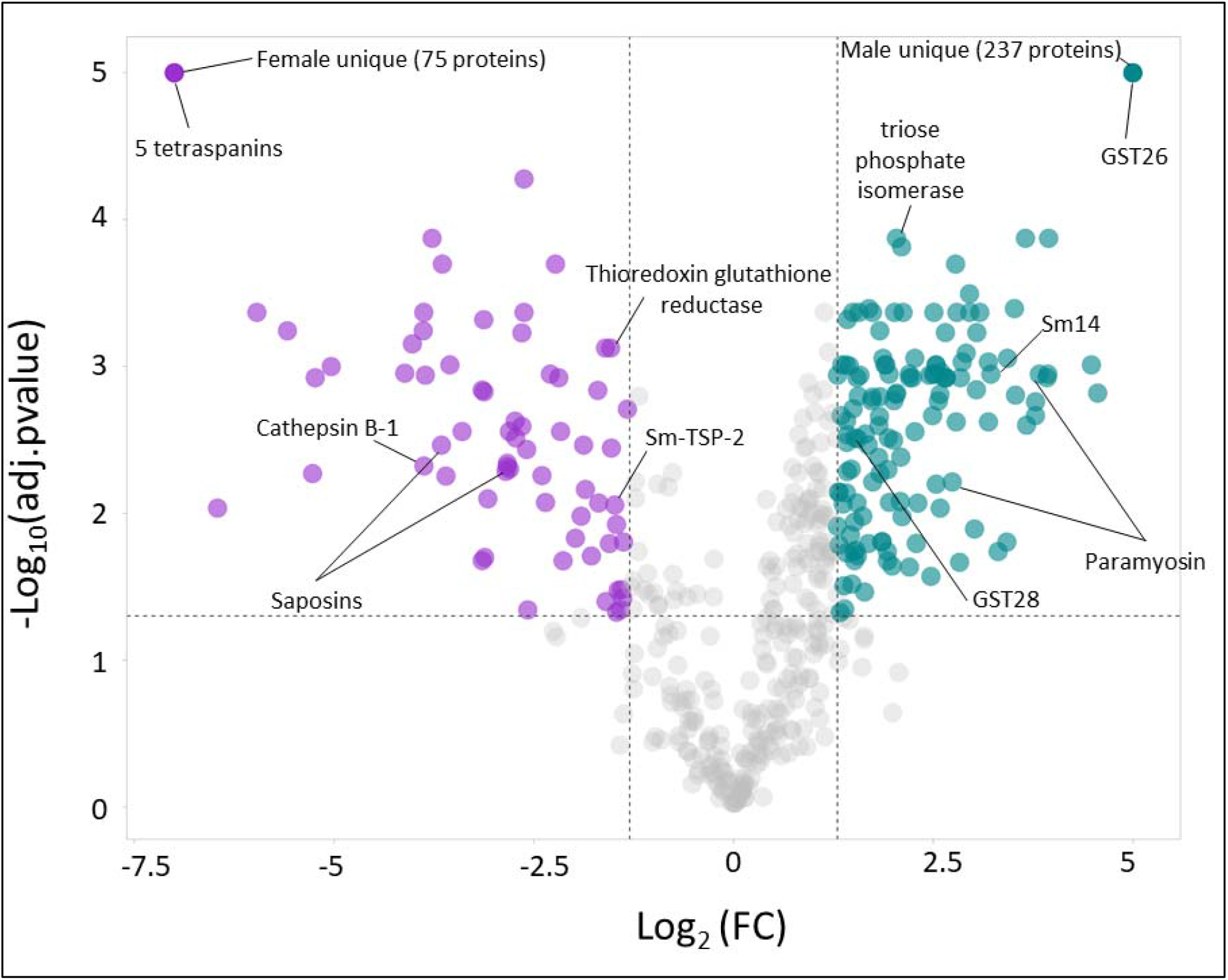
Volcano plot highlighting the proteins with significantly differential relative protein abundance levels in the secretomes of male (M) and females (F) *Schistosoma mansoni* flukes.

When comparing MF vs. M, 27 and 15 proteins were exclusively identified in MF and M, respectively and could not be, thus, quantified. However, the relative abundance of the remaining 700 proteins was not significantly different (Supplementary Figure 3), suggesting that males have a bigger contribution to the total MF secretome than females. It is worth noting that some well-characterised Schistosoma spp. vaccine candidates (26) were upregulated or uniquely found in the secretome of females (i.e., A0A5K4F8N6_SCHMA (Sm-TSP-2), Q8MNY2_SCHMA (cathepsin B-1), A0A5K4EE66_SCHMA (Thioredoxin glutathione reductase) or males (i.e. A0A3Q0KIP4_SCHMA, A0A3Q0KD88_SCHMA (paramyosin), FABP_SCHMA (Sm14), GST26_SCHMA (Glutathione S-transferase 26), G4V6B9_SCHMA (triose phosphate isomerase)). Furthermore, two saposins (G4VBU4_SCHMA and A0A3Q0KK40_SCHMA) were also upregulated in the secretome of females. It is also worth highlighting that five different tetraspanins (A0A3Q0KTH5_SCHMA, G4LUN6_SCHMA, A0A5K4FD80_SCHMA, Q86D97_SCHMA and G4LWW2_SCHMA) were uniquely found in the secretome of females, while Sm-TSP-2 was found upregulated in the secretome of females.

### 3.2 *S. mansoni* male adult worms secrete proteins implicated in carbohydrate metabolism, redox functions and cytoskeletal organisation

A protein-protein association (PPA) analysis of proteins uniquely secreted or upregulated in the secretome of *S. mansoni* adult males revealed a strong network of interacting proteins. Approximately 70% of the proteins interacted in one or multiple clusters with other proteins. While most unique and upregulated proteins interacted together, one cluster of male-unique proteins could be differentiated (Fig. 2A, circle), which included splicing factors, RNA binding proteins and ribonucleoproteins. Furthermore, a clear cluster of proteins with upregulated abundance in males was found to be associated with glycolysis and gluconeogenesis pathways (Fig. 2A, rectangle).

**Figure 2.**
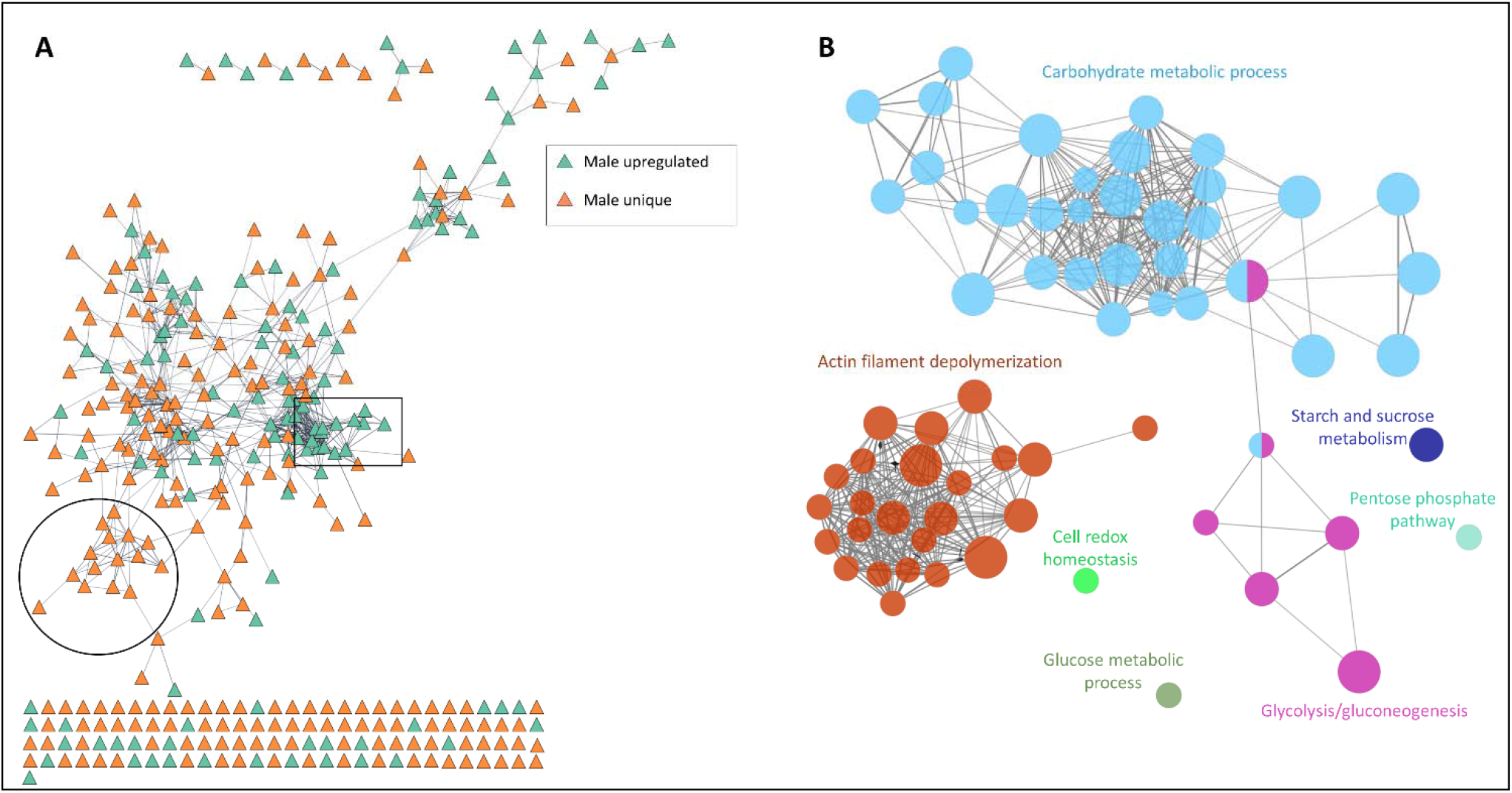
Functional analysis network of proteins uniquely secreted or upregulated in the secretome of adult *Schistosoma mansoni* males. (A) Protein-protein association network of all proteins uniquely present or with significantly higher abundance in the secretome of adult *S. mansoni* males. (B) Functional analysis network showing the gene ontology terms and KEGG pathways with the highest significance.

The functional analysis showed at least 7 groups (adjusted P-value < 0.0005) containing 67 non-redundant biological terms (adjusted P-value < 0.05), including “glucose metabolic process” (GO:0006006), “cell redox homeostasis” (GO:0045454), “pentose phosphate pathway” (KEGG:00030), “starch and sucrose metabolism” (KEGG:00500, “glycolysis/Gluconeogenesis (KEGG:00010), “actin filament depolymerization” (GO:0030042) and “carbohydrate metabolic process” (GO:0005975) (Fig. 2B, Supplementary Table 3).

The cluster containing more terms (31 nodes) and proteins (total of 54 proteins) was associated with pathways involved in carbohydrate metabolism (a gene ontology term also associated with the KEGG pathway glycolysis/gluconeogenesis) (Fig. 2B, Supplementary Table 3). Proteins involved in this pathway included phosphoenolpyruvate carboxykinase, glucose-6-phosphate dehydrogenase and malate dehydrogenase among others. The second cluster with the highest number of terms (24 nodes) and proteins (22 proteins) was associated with a cytoskeleton organisation function, and included proteins such as calponin, paramyosin, filamin, and tubulin.

### 3.3 *S. mansoni* female adult worms secrete proteins with hydrolase activity and cellular homeostasis

The PPI analysis of proteins uniquely secreted or upregulated in the secretome of *S. mansoni* adult females showed that only around 50% of the proteins interacted in one or multiple clusters with other proteins. Contrary to that observed for male schistosomes, uniquely female-secreted proteins or proteins with an upregulated expression in the secretome of females did not cluster together.

The functional analysis showed at least 6 groups (adjusted P-value < 0.005) containing 54 non-redundant biological terms (adjusted P-value < 0.05), including “lysosome” (KEGG:04142), “calcium-dependent phospholipid binding” (GO:0005544), “transmembrane transporter activity” (GO:0022857), “hydrolase activity” (GO:0016798), “peptidase activity” (GO:0070011) and “transition metal ion transport” ((GO:0000041) (Fig. 3B, Supplementary Table 4). The cluster with the highest number of terms and proteins was associated with the cellular homeostasis process (28 nodes and 20 proteins) and included proteins such as ferritin and thioredoxin glutathione reductase (Fig. 3B, Supplementary Table 4). Additionally, two clusters associated with peptidase activity (9 nodes and 15 proteins) and hydrolase activity (7 nodes and 11 proteins) were also highlighted to be of importance in this analysis (Fig. 3B, Supplementary Table 4). These two clusters included proteins such as galactosidase, glucosidase, cathepsin B, cathepsin L, carboxypeptidase and other cysteine- and serine-peptidases (Supplementary Table 4).

**Figure 3.**
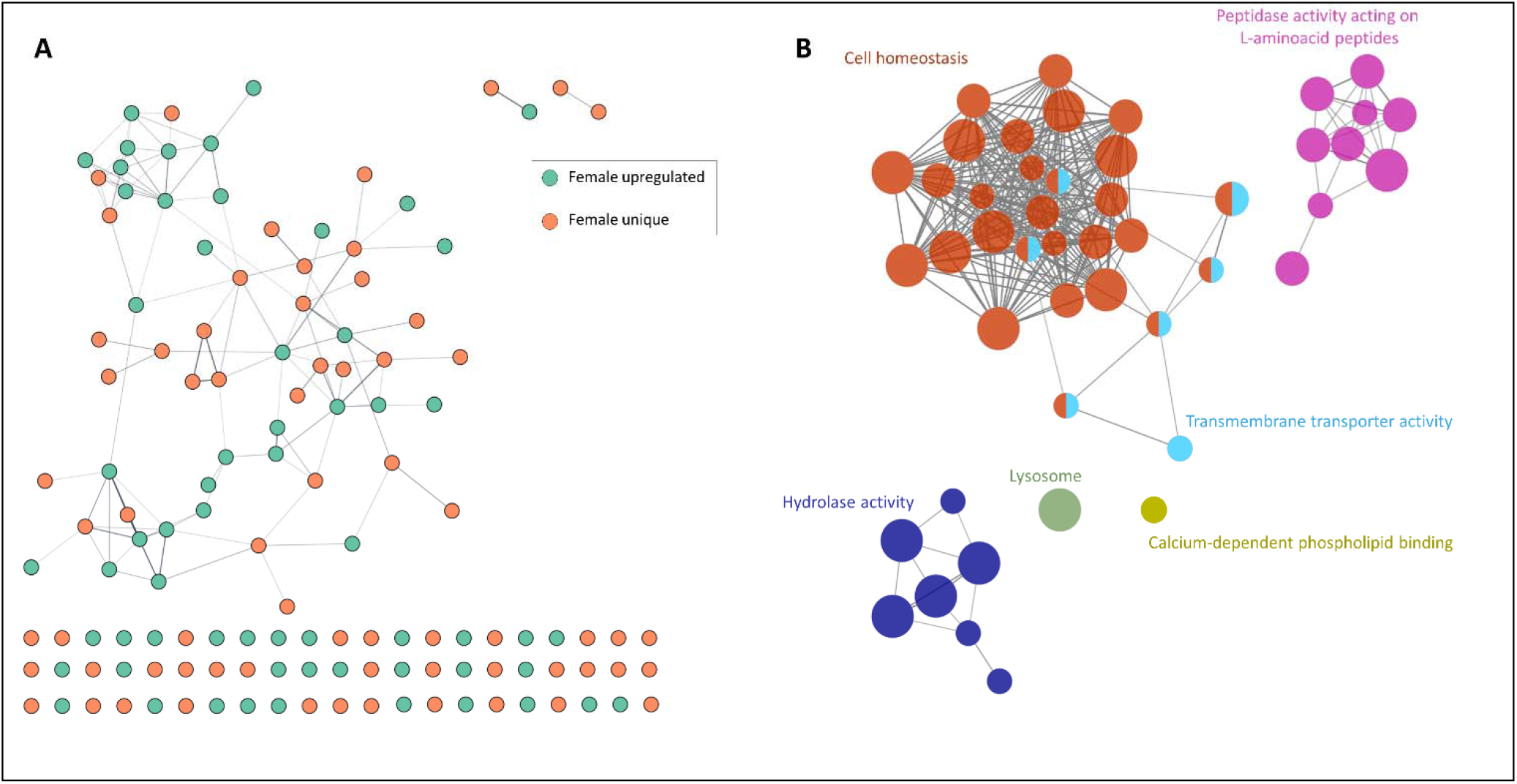
Functional analysis network of proteins uniquely secreted or upregulated in the secretome of *Schistosoma mansoni* adult females. (A) Protein-protein association network of all proteins uniquely present or with a significantly upregulated expression in the secretome of *S. mansoni* adult females. (B) Functional analysis network showing the gene ontology terms and KEGG pathways with the highest significance.

## 4 Discussion

Schistosomes are one of the most important groups of human helminths in terms of public health. Unlike other trematodes, they are usually found as paired couples, with the female adult worm living within the gynaecophoric canal of the male (McManus et al., 2018). Indeed, this dioecy distinguishes schistosomes from other flatworms that infect humans. This sexual dimorphism has different biological implications, including the need for pairing to achieve sexual maturity in females (reviewed in (Moore et al., 1954)), and favouring a division of labour between the more muscular male migrating towards oviposition sites and the filiform female reaching small vessels to discharge the eggs (Loker and Brant, 2006). Yet despite their importance in male-female communication, only a handful of studies have analysed in depth the molecules and receptors involved in these interactions (Armstrong, 1965;Basch and Basch, 1984;Gupta and Basch, 1987;Chen et al., 2022). In this study we aimed to comprehensively characterise the protein complement of the ESPs from adult stage female and male *S. mansoni* to increase knowledge into the biology of these worms and to augment the repertoire of potential diagnostic and vaccine candidates for schistosomiasis.

Molecules released and secreted by schistosomes, including tegumental proteins and digestive enzymes (in the vomitus (Hall et al., 2011)) play a key role in host-parasite and male-female interactions, and their expression and secretion is driven by the divergent requirements and functions of both schistosome sexes. For instance, in *S. japonicum* and *S. bovis*, the male schistosome exhibits significantly more proteins in its tegument than the female, both in total number and in unique proteins(Perez-Sanchez et al., 2008;Zhang et al., 2013)). Our present findings are in agreement with this situation, and indeed are not unexpected given the larger size of the male schistosome. Furthermore, the protein and small RNA content of *S. japonicum*-secreted extracellular vesicles and the expression of phosphoproteins from *S. mekongi* also are sex-dependant (Du et al., 2020). Our results also revealed a gender-specific divergence in the secretome landscape in *S. mansoni*, which, as shown at the transcriptomic level, could be beneficial for the sexual dimorphism in this species (Fitzpatrick et al., 2005;Anderson et al., 2015;Picard et al., 2016). Recent studies have shown that *S. japonicum* female-tegumental proteins are involved in protein glycosylation and lysosome function, while male-tegumental proteins play a role in intracellular signal transduction, regulation of actin filament polymerization, and proteasome core complex (Zhang et al., 2013). Our results also showed functions related to actin filament depolymerisation and lysosome to be important in male and female-secreted proteins, respectively. While proteins belonging to the lysosome KEGG ontology is a markedly heterogeneous group including cathepsin B peptidases, tetraspanins and glycosidases, proteins related to actin filament depolymerisation play a specific role in cytoskeletal regulation. These results confirm previous transcriptomic studies and support the hypothesis of the role of the male schistosome in physical support of the female to facilitate migration against the flow of the portal circulation toward the mesenteric venules where the female schistosome deposits the eggs (Fitzpatrick et al., 2005;Cai et al., 2016;Phuphisut et al., 2018).

Glucose metabolism is essential in female worms due to the energy requirement to support the production and release of the large number of eggs laid daily - around 300 eggs per female per day in *S. mansoni* (Cheever et al., 1994). Glycogen consumption, however is paramount in other schistosome functions such as muscle contraction and tegumental membrane repair, both being significantly enriched among proteins more abundant in the adult male versus the female schistosome (Gobert et al., 2003). Earlier transcriptomic investigations revealed that the expression of glucose transporters including gtp1 and gtp4 was not influenced by the sex of the schistosome (Cai et al., 2016). By contrast, we observed significantly elevated abundance of GTP1 (Smp_012440, Q26579_SCHMA) and of other glucose transporters (Smp_105410, G4VC44_SCHMA) in the secretome of the female *S. mansoni*, likely the result of an increased expression in the tegument. These discrepancies could be in part explained since transcriptomic analysis was performed on whole worms whereas our study focused solely on the ESPs. Furthermore, other key glycolytic enzymes such as aldolase (Smp_042160.1, ALF_SCHMA) and glycerol-3-phosphate dehydrogenase (G3PDH, Smp_030500.1, C4Q5J8_SCHMA) were more abundantly secreted by males. Our results agreed with previous findings that reported the higher consumption of glucose and importance of glycogen storage in the male worm, which could reflect the muscular effort involved in transporting the female through the portal vasculature, and the need by the female to transport these molecules from the male (Skelly et al., 2014). Notably from the viewpoint of infection control, aldolase and G3PDH have been a focus of vaccines and other intervention targets (Dessein et al., 1988;Goudot-Crozel et al., 1989;Tallima et al., 2017).

In addition, the findings indicate that enzymes such as hydrolases and peptidases play an important role in the biology of female schistosomes. Hydrolases are a common group of proteins that include lipases, phosphatases, and glycosidases among others. In the case of female schistosomes, several glycosidases were found more abundantly secreted, including beta-glucosidase (Smp_043390, A0A3Q0KF32_SCHMA), alpha-galactosidase (Smp_089290, G4VLE3_SCHMA), and several alpha-amylases. Furthermore, it has been suggested that hydrolases released by the schistosome egg contribute to the transit of the egg and the circumoval granuloma across the intestinal wall and also with the nutritional requirements of the embryo (Cesari et al., 2000). Despite our findings, we cannot rule out that hydrolases detected here could have been secreted by eggs and did not strictly originate from the adult female. Other peptidases including cathepsins participate in invasion of the skin by the cercaria (Dvorak et al., 2008), haemoglobin degradation by the adult stage (Gotz and Klinkert, 1993;Dalton et al., 1995), and egg hatching (Rinaldi et al., 2009), and have prominent immunogenic properties (Soloviova et al., 2019). Females secreted several cathepsins more abundantly than males, including SmCB1 (Q8MNY2_SCHMA), which is known to be secreted from the gut of schistosomes and to contribute to Th2 polarization responses (de Oliveira Fraga et al., 2010). This enzyme has been validated as a potent anti-schistosome chemotherapeutic target (Abdulla et al., 2007;Jilkova et al., 2021).

We found a number of other well-characterised vaccine candidates upregulated in the secretome of male worms (i.e., Sm14 (Smp_095360.1, FABP_SCHMA), GST26 (Smp_163610.1, GST26_SCHMA), GST28 (Smp_054160.1, GST28_SCHMA), three paramyosin isoforms (Smp_046060.1, A0A3Q0KFC2_SCHMA; Smp_085540.6, A0A3Q0KIP4_SCHMA; Smp_021920.3, A0A3Q0KD88_SCHMA)) as well as in the secretome of females (i.e. Sm-TSP-2 (Smp_335630.1, A0A5K4F8N6_SCHMA), thioredoxin glutathione reductase (A0A5K4EE66_SCHMA), cathepsin B-1 (Smp_103610.1, Q8MNY2_SCHMA) and several saposins (Smp_194910.1, G4VHH1_SCHMA; Smp_130100.1, G4VBU4_SCHMA, Smp_105450.1, A0A3Q0KK40_SCHMA)). Unexpectedly, most tetraspanins identified were significantly upregulated in the secretome of female worms. Sm-TSP-2 formulated with glucopyranosyl lipid adjuvant has proven safe in a phase I trial (Keitel et al., 2019), and a homolog in *S. haematobium* (Sh-TSP-2) showed efficacy in a heterologous mouse model of schistosomiasis (Mekonnen et al., 2020). Furthermore, tetraspanins have been successfully tested as diagnostic candidates against urogenital schistosomiasis (Pearson et al., 2021;Mekonnen et al., 2022). Interestingly, we found five other tetraspanins uniquely present in the secretome of females, and their study as vaccine or diagnostic candidates could be of interest. It is noteworthy that both GSTs (GST26 and GST28) were upregulated in the secretome of males. Although GSTs have been widely studied as vaccine candidates (Riveau et al., 1998), recent reports reveal a lack of efficacy against urogenital schistosomiasis in children (Riveau et al., 2018).

We have additionally identified 75 and 237 proteins uniquely secreted by female and male worms, respectively. These proteins likely play key roles in schistosome biology, notably male-female communication. Recent reports have highlighted the secretion by male schistosomes of a specific small molecule (ß-alanyl-tryptamine) that is key for the development and laying of eggs by females (Chen et al., 2022) (ß -alanyl-tryptamine is a small peptide of ∼300 Daltons in mass which would not have been retained when our samples were prepared during centrifugation which employed a 3 kDa cutoff membrane.) Furthermore, in *S. japonicum*, biogenic amine neurotransmitters have been shown to be also highly implicated in male-female sexual communication (Wang et al., 2017). Based on the present findings, we posit that the development of drugs interrupting male-female communication could lead to novel and effective control measures.

To conclude, a better breadth of coverage of the adult stages of *S. mansoni* ESP profile, and a deeper understanding of the most highly secreted proteins will be of importance for basic science aimed at understanding schistosome biology, thus will provide important information for the development of novel vaccine strategies against this major neglected tropical disease. Moreover, identification of the most abundantly secreted proteins of both sexes enables future analysis of the regulatory elements and motifs that control the expression of the corresponding genes can assist with the development of transgenic schistosomes that over-express endogenous proteins, or even secrete foreign proteins. Access by the field to transgenic schistosomes that (conditionally) secrete reporters, model antigens, and other informative gene products, along with advances in human challenge models (Langenberg et al., 2020) can be expected to lead to noteworthy progress in the immunobiology and pharmacology of these flukes (Hoffmann et al., 2014;Zamanian and Andersen, 2016;McVeigh and Maule, 2019;Douglas et al., 2021;Quinzo et al., 2022).

## 7 Author Contributions

VHM, ETK, WI, PJB and JS designed the experiments, ETK, MM, BAR, BKB, AL, PJB and JS analysed the data, and JS, ETK, VHM, and PJB drafted the manuscript with input from all the co-authors; WI, EKT, VHM contributed the helminth materials JS performed mass spectrometry focused analysis; JS, AL, and PJB supervised the project. JS, AL, MM, PJB and BKB arranged the funding. All authors read and approved the final draft.

## 9 Acknowledgments

The proteomic analysis was performed in the proteomics facility of SCSIE University of Valencia. For the purposes of Open Access, the authors have applied a CC BY public copyright license to any Author Accepted Manuscript version arising from this submission. Schistosome-infected mice and adult schistosomes were provided by the NIAID Schistosomiasis Resource Center of the Biomedical Research Institute, Rockville, Maryland through NIH-NIAID Contract HHSN272201000005I for distribution through BEI Resources. The findings were obtained from work partially supported by the Defense Advanced Research Projects Agency (DARPA) and Naval Information Warfare Center Pacific (NIWC Pacific), under Contract No. N66001-21-C-4013 (Approved for Public Release, Distribution Unlimited). The views, opinions, and/or findings expressed are those of the authors and should not be interpreted as representing the official views or policies of the Department of Defense or the U.S. Government.

## 10 Data Availability Statement

Mass spectrometry data along with the identification results have been deposited in the ProteomeXchange Consortium via the PRIDE partner repository with the dataset identifier PXD030699.

## 11 Figure legends

**Supplementary Figure 1**. Data normalisation using medians of summed intensities after label-fere quantitative analysis.

**Supplementary Figure 2**. Power calculation and fasle discovery rate of the label-free quantitative analysis performed using MSstats.

**Supplementary Figure 3**. Volcano plot of *Schistosoma mansoni* secreted proteins from male vs male-female (mix). No statistically significant differences were observed.

