## Supplementary figures and images for "Differential excretory/secretory proteome of the adult female and male stages of the human blood fluke, *Schistosoma mansoni*"

### Supplementary Figure 1

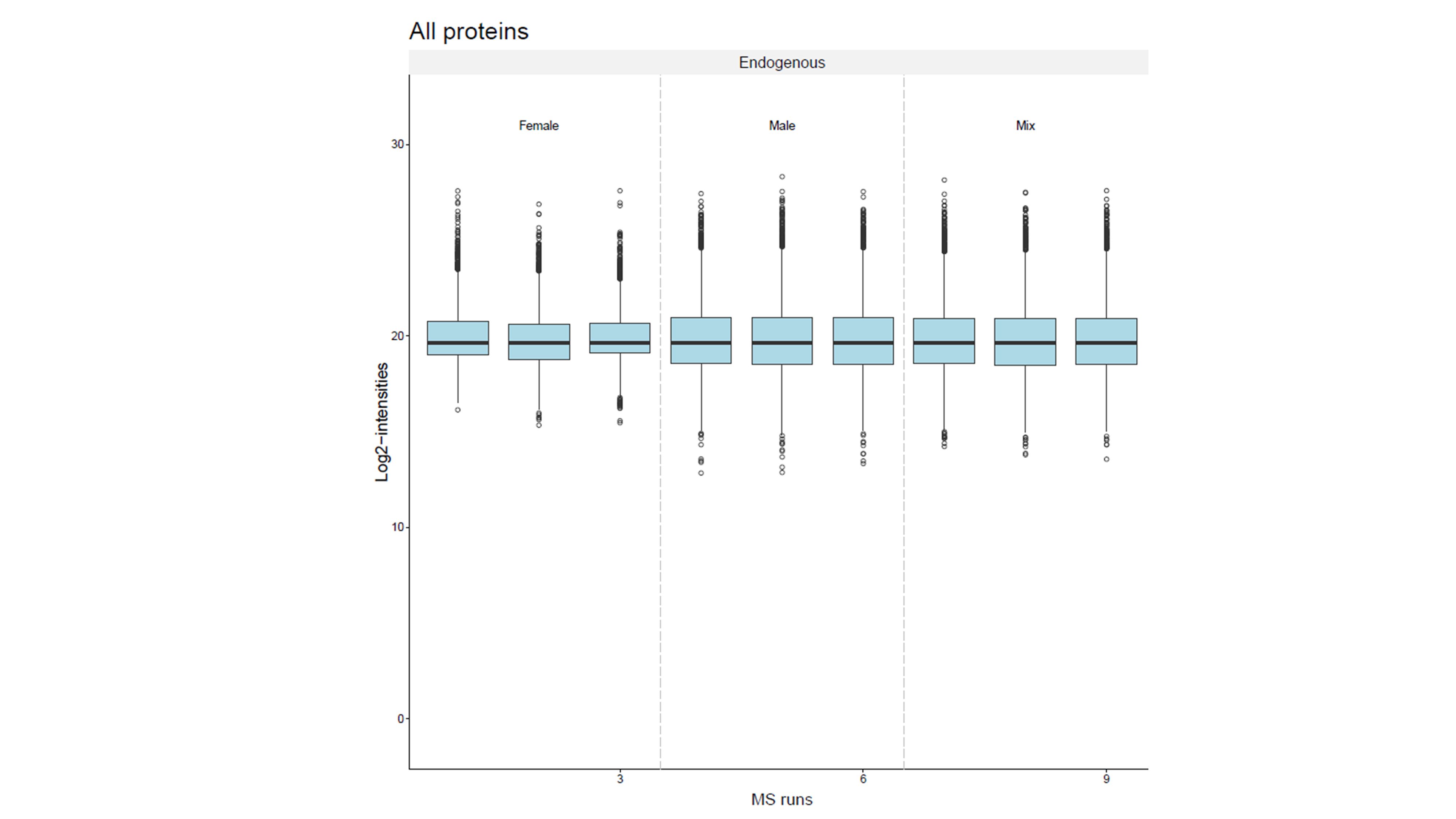

### Supplementary Figure 2

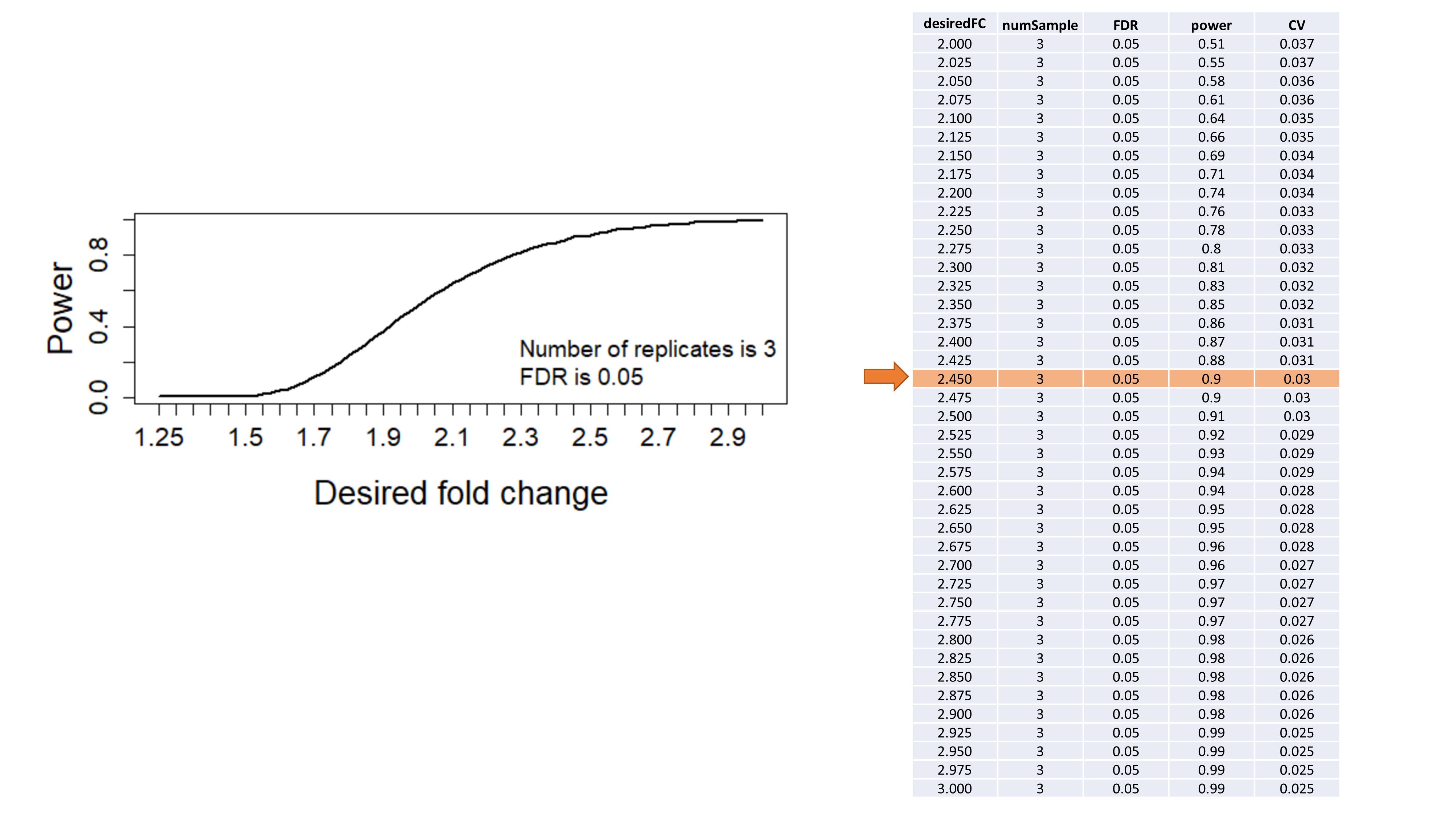

### Supplementary Figure 3

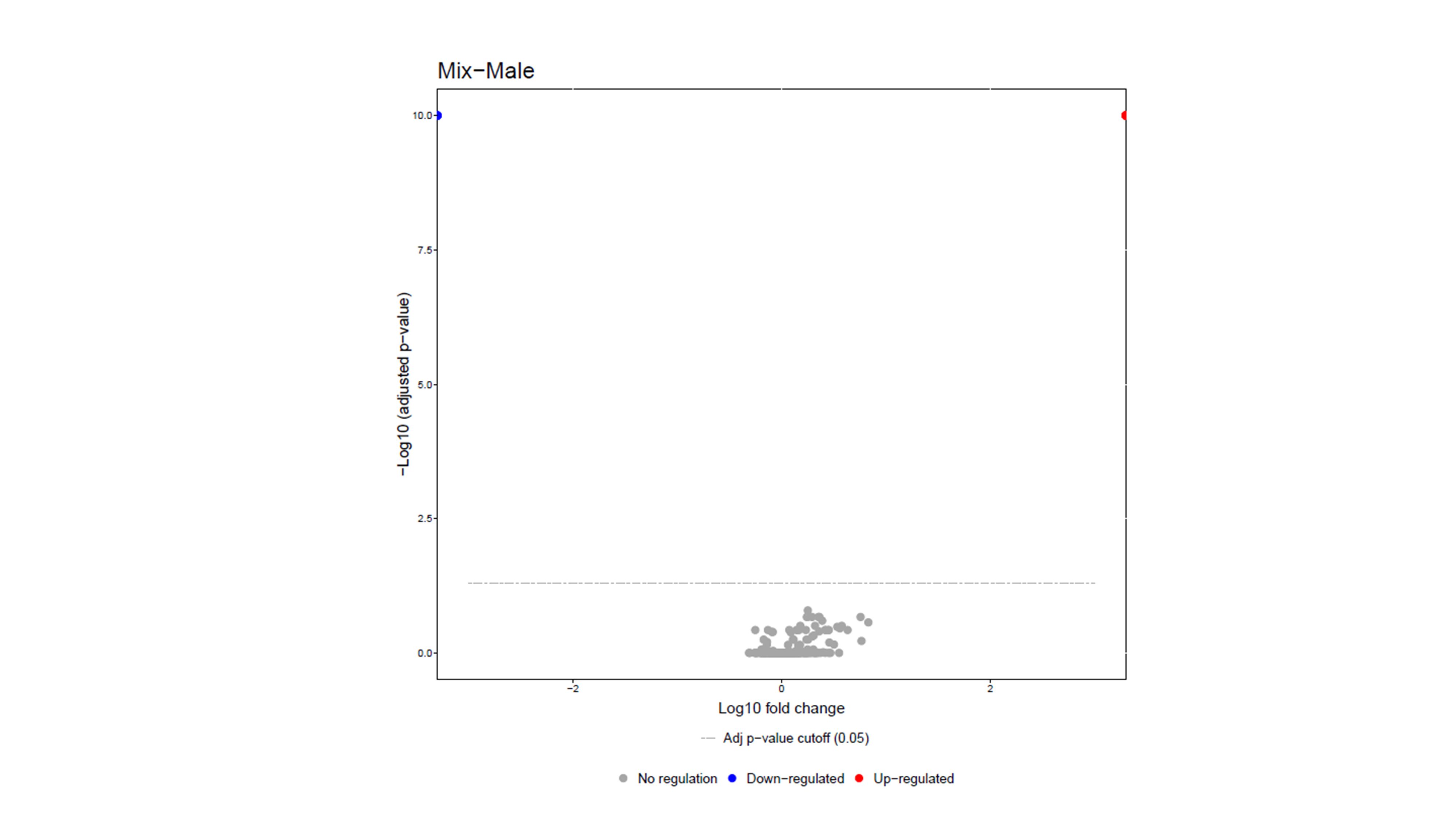
